# Motor variability prior to learning is a poor predictor of the ability to adopt new movement solutions

**DOI:** 10.1101/2020.10.23.350819

**Authors:** Rajiv Ranganathan, Marco Lin, Samuel Carey, Rakshith Lokesh, Mei-Hua Lee, Chandramouli Krishnan

## Abstract

Many contexts in motor learning require a learner to change from an existing movement solution to a novel movement solution to perform the same task. Recent evidence has pointed to motor variability prior to learning as a potential marker for predicting individual differences in motor learning. However, it is not known if this variability is predictive of the ability to adopt a new movement solution for the same task. Here, we examined this question in the context of a redundant precision task requiring control of motor variability. Fifty young adults learned a precision task that involved throwing a virtual puck toward a target using both hands. Because the speed of the puck depended on the sum of speeds of both hands, this task could be achieved using multiple solutions. Participants initially performed a baseline task where there was no constraint on the movement solution, and then performed a novel task where they were constrained to adopt a specific movement solution requiring asymmetric left and right hand speeds. Results showed that participants were able to learn the new solution, and this change was associated with changes in both the amount and structure of variability. However, individual differences in baseline motor variability were only weakly correlated with initial and final task performance when using the new solution, with greater variability being associated with higher errors. We also found a strong specificity component – initial variability when using the new solution was highly correlated with final task performance with the new solution, but once again, higher variability was associated with greater errors. These results suggest that motor variability is not necessarily indicative of flexibility and highlight the need to consider the task context in determining the relation between motor variability and learning.

---

The presence of motor redundancy at several levels in the human body and the task gives rise to the phenomenon that a given task goal can be achieved using multiple movement solutions. Although the evidence for how participants exploit this redundancy in the body and the task has been well described (Cohen and Sternad 2009; Cusumano and Cesari 2006; Ranganathan et al. 2013; Scholz and Schöner 1999; Todorov and Jordan 2002), the question of how ‘flexible’ participants are in changing from one movement solution to another is less well understood. Understanding individual differences in such flexibility is critical since several contexts such as coaching and neurorehabilitation require participants to learn a new movement solution to perform the same task.

One potential feature that is relevant to predicting how flexible an individual is in adopting a new movement solution is motor variability. In task with redundancy, variability can be split into two components (Cusumano and Cesari 2006; Mosier et al. 2005; Scholz and Schöner 1999) – (i) ‘task space’ variability, i.e., the component of movement variability that affects task outcome, and (ii) ‘null space’ variability, i.e., the component of movement variability that does not affect the task outcome. Because flexibility is a measure of how well participants can move from one point in the null space to another, the null space variability has been hypothesized as a measure of how flexible participants are, with greater null space variability (relative to the task space variability) indicative of stronger synergies (Latash et al. 2002). More recently, motor variability has also been shown to predict individual differences in motor learning – when participants have greater variability, they can engage in more efficient exploration strategies to facilitate learning (Wu et al. 2014). Although the generality of this finding has been questioned (He et al. 2016; Singh et al. 2016), overall these results support the view that motor variability is not simply noise (Newell and Corcos 1993), and can be a potential signature for predicting future motor learning (Dhawale et al. 2017).

However, in understanding the role of motor variability in predicting individual differences in learning, two issues remain unaddressed. First, the issue of flexibility (i.e., how well participants can move from one solution to another) has received little attention. A critical component for understanding this question is the use of tasks that have redundancy. Second, prior studies on using variability to predict individual differences in motor learning have primarily focused on adaptation tasks (He et al. 2016; Singh et al. 2016; Wu et al. 2014). Although adaptation is one form of motor learning, a more common form of motor learning relevant to real-world contexts is skill learning, where there is a relatively permanent change in the underlying movement capacity (Krakauer and Mazzoni 2011; Schmidt and Lee 2011; Sternad 2018). One such class of skill learning tasks are precision tasks that require learning to produce a consistent outcome over multiple trials (Cohen and Sternad 2009; Muller and Sternad 2004; Ranganathan and Newell 2010a; Shmuelof et al. 2012). Here, we address both these issues using a precision task that has redundancy.

The goals of this study were to (i) characterize changes in task performance and movement variability when learning a new solution to perform a task, and (ii) examine if motor variability prior to learning is predictive of motor performance when learning a new solution to a redundant motor task. Participants learned a bimanual throwing task which required participants to slide a virtual puck to a specified target. The motion of the puck depended on the sum of the speeds of the two hands, making the task redundant. We exploited the fact that participants in bimanual tasks tend to typically use symmetric (or near-symmetric) contributions from both hands (Kelso et al. 1979; MacKenzie and Marteniuk 1985), and designed a task that required them to switch to a novel asymmetric solution to this task. We anticipated that if motor variability is predictive of individual differences in learning new solutions, then the motor variability observed at baseline (i.e., prior to the learning of the new solution) should be predictive of performance using the new solution, with higher variability being associated with better task performance (i.e. lower errors).

## Methods

### Participants

Participants were college-aged adults with no history of movement impairments in the upper extremity (N = 50, age 18-25 years, 40 women) and were naive to the purpose of the experiment. All participants provided written consent and the experimental protocol was approved by the Michigan State University Institutional Review Board.

### Apparatus

The participants performed all the tasks on a two joint bimanual end-point robot (KINARM technologies, Kingston, ON) (Fig. 1a). The position data from the two robot handles were sampled at 1000 Hz. The visual display was set up through a semi-silvered screen so that images were shown in the plane of the hands and direct vision of the hands was obstructed.

**Figure 1.**
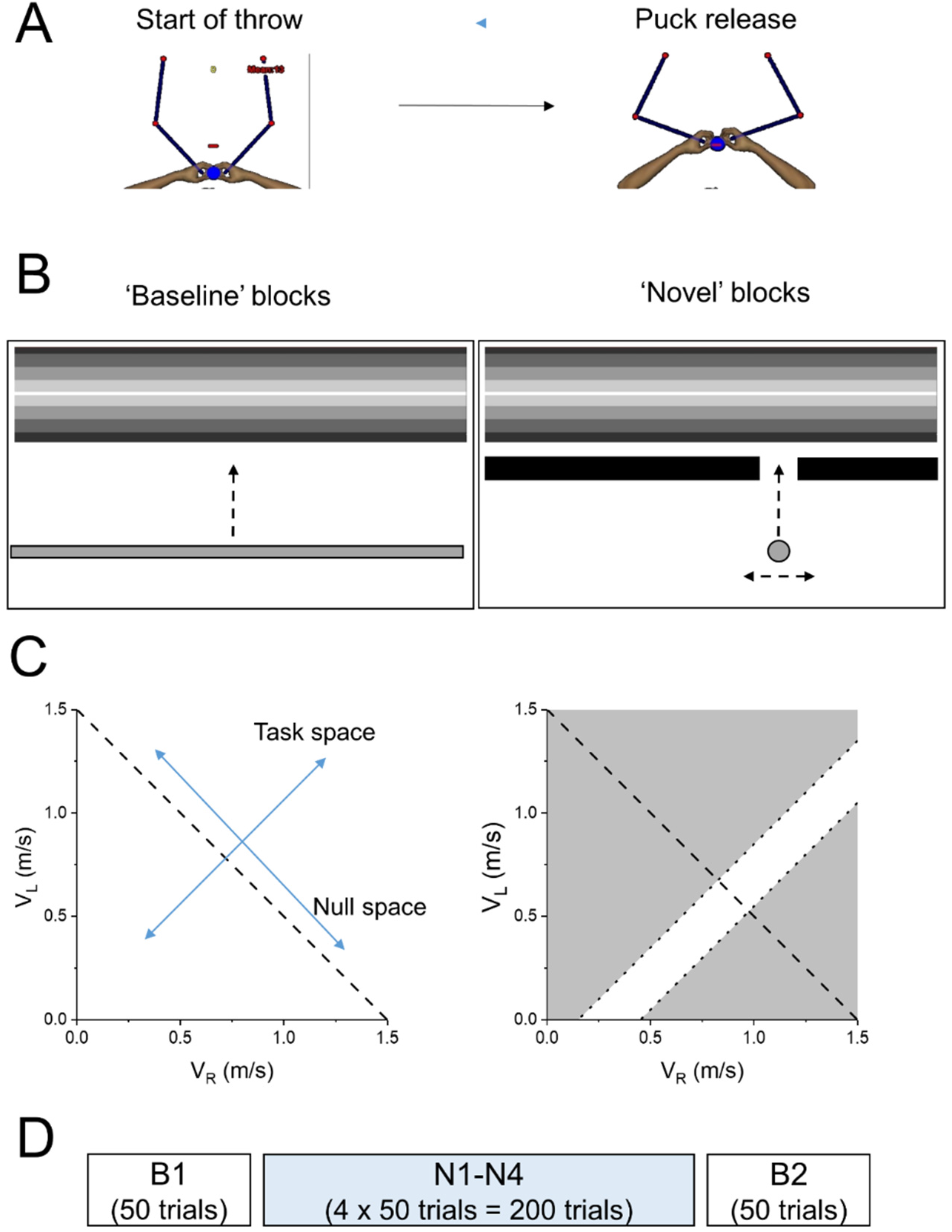
Schematic of task and experiment. (A) Virtual shuffleboard task – participants grasped a bimanual robot and performed a throwing motion toward a slot located 10 cm away. When the puck crossed this slot, the puck was ‘released’ and participants saw visual feedback indicating their task performance (B) Visual feedback of task performance. During the baseline blocks, participants saw the puck as a ‘horizontal’ bar and a puck speed of 1.5 m/s would land the bar right on the middle of the target (indicated in white). During the novel blocks, participants saw the puck as a circle. The vertical motion of the puck was exactly the same as the horizontal bar in the baseline blocks, but the horizontal motion of the puck was determined by the difference in the right and left hand speeds. Participants had to select specific solutions to make sure that the ball passed through the ‘hole’ in a wall. (C) Solution space in terms of left and right hand speeds. In the baseline blocks (left), the solution manifold is represented by the dotted line, which indicates the speed combinations that lead to a total of 1.5 m/s. In the novel blocks (right), the presence of the wall reduces the solution manifold to a much smaller area (areas shaded in grey would lead to a collision of the puck with the wall). (D) Experimental protocol. After an initial familiarization, all participants completed a baseline block (B1), followed by four novel blocks (N1-N4), and a return to the baseline block (B2).

### Task

The task used was a virtual shuffleboard task where the goal of the participant was to slide a virtual puck toward a target shown on the screen (Cardis et al. 2018) (Fig. 1a). At the start of each throw, participants were instructed to position both hands in a respective ‘home’ position. At this point, the individual hand cursors disappeared and were replaced by a circular puck at the average position of the hands. They were then asked to slide the puck toward a slot positioned straight ahead. Once participants crossed the slot, the puck was ‘released’ and a second screen was shown where the puck traveled towards a target. The speed of the puck was dependent on the sum of the speeds of the left and right hands at release (i.e., V_puck_ = V_L_ + V_R_) and perfect task performance was achieved when V_puck_ = 1.5 m/s. Because the speed is dependent on both hands, multiple solutions can be used to achieve perfect task performance.

There were two versions of the task (Fig. 1b). In the ‘baseline’ version of the task, participants were not constrained to use any specific solutions for the task. Visual feedback of the puck was provided as a horizontal line stretching across the screen. After each throw, participants saw the position of the horizontal line and were provided a numerical score depending on the error (with a max of 100 points). In the ‘novel’ version of the task, participants were constrained to use a specific set of solutions. Visual feedback was provided as a circular puck. The vertical motion of the puck was identical to the motion of the line in the baseline blocks, and the horizontal dimension was controlled by the difference in the hand velocities so that a higher velocity on the right (left) hand would move the puck further to the right (left). We then constrained the solution adopted by adding a wall with a hole in a specific region. The hole was placed on the right-hand side of the screen so that participants now had to produce a higher velocity on the right hand by 0.3 ± 0.15 m/s to make the puck pass through the hole. Therefore, for the puck to pass through the center of the hole and land perfectly on the target, the combination would be (V_R_, V_L_) = (0.9, 0.6) m/s – i.e., the V_L_+V_R_ needs to be 1.5 m/s and V_R_ would have to be 60% of the total speed. The scoring system was the same as the baseline blocks with the addition that if the puck collided against the wall, that trial was scored as zero points. The solution space in terms of the two-hand speeds for both the baseline and novel blocks is shown (Fig. 1c)

### Procedure

Participants were first given 10 familiarization trials in each condition to make sure they understood the goal of the task. This was followed by 6 blocks of practice – a baseline block (B1), 4 novel blocks (N1-N4), and a baseline block (B2) (Fig. 1d). Each block consisted of 50 throws and the entire experiment was done in a single session that lasted about 45 minutes.

## Data Analysis

### Task performance

#### Absolute error

Because perfect task performance was achieved when the puck release speed was 1.5 m/s, we computed the absolute error as the absolute difference between the actual puck speed at release and the desired release speed of 1.5 m/s.

#### Collisions

To indicate how well participants adapted to the novel condition, we computed the frequency of trials in which the ball collided with the wall in each block (relative to the total number of trials in the block). A higher number of collisions indicated greater difficulty with changing to the novel condition. Because there was no wall in the baseline conditions, the number of collisions in the baseline blocks is zero by definition.

### Coordination

#### Speed ratio

The coordination in this task was measured as the contribution of the right hand to the puck speed – i.e., V_R_/(V_R_+V_L_). As mentioned earlier, in the baseline blocks, there was no constraint on the coordination, but a symmetric contribution from both hands would result in a ratio of close to 0.5. During the novel blocks, given the position of the obstacle, this ratio had to be close to 0.6 for the puck to pass through the center of the hole.

### Variability measures

#### Movement variability

In each block, we split the movement variability along two orthogonal dimensions-the task space and the null space (as shown in Fig. 1c). The variability along each of these dimensions is referred to as the task and null space variability (Cardis et al. 2018)

#### Autocorrelation

In addition to the amount of variability, we also computed the lag-1 autocorrelation in the task and null spaces as a measure of the structure of variability (Abe and Sternad 2013; van Beers et al. 2013; Cardis et al. 2018; Dingwell et al. 2013). Each trial, represented by a point in the (V_L_, V_R_) space was projected on to the task space and the null space and the time series of these two projections was used for the autocorrelation analyses. A positive lag-1 autocorrelation indicates that deviations tend to persist over time, whereas a negative lag-1 autocorrelation indicates that a positive deviation is more likely to be followed by a negative deviation on the next trial.

## Statistical Analysis

### Learning a new solution

We first quantified the learning in the task by observing changes over the practice blocks. For all dependent variables except collisions, this was analyzed by a one-way ANOVA with block as the main factor (6 levels- B1, N1, N2, N3, N4 and B2). We performed three a priori comparisons related to this ANOVA by comparing the following blocks: (i) B1 and N1 (i.e., what happens when participants initially switch to the new solution), (ii) N1 and N4 (i.e., what happens when they learn the new solution), and (iii) N4 and B2 (i.e., what happens when they change from new solution back to baseline). Corrections for violation of sphericity were performed using the Greenhouse-Geisser correction when appropriate. For the analysis of collisions, which only occurred in the novel blocks, we used a paired t-test to compare the collisions in N1 and N4. The level of significance was set at .017 (corrected for the 3 comparisons). All analyses were run in JASP version 0.9 (JASP Team 2018).

### Predicting individual differences

The primary focus was to examine if the initial and final performance on the novel task could be predicted by motor variability at baseline. So we examined Pearson’s correlations between: (i) absolute error in N1 with task space and null space variability in B1, and (ii) absolute error in N4 with task and null space variability in B1. In addition, we also computed if the final performance on the novel task could be predicted using the variability on the first block of the novel task and computed the correlation between absolute error in N4 with the task and null space variability in N1. Since we examined a total of six correlations, we chose a Bonferroni correction so that the significance threshold was set at p = .0083. Since there was a pair of correlation coefficients (one correlation with the task space variability and one with the null space variability), we also compared these two correlation coefficients using the cocor package (Diedenhofen and Musch 2015) to examine if the correlation in one space was bigger than the other.

## Results

Based on Tukey’s boxplots of the absolute error in B1, data from two participants were excluded from analysis. Both individuals had higher error in B1 than the Tukey’s criterion.

### Learning a new solution

Participants showed an initial increase in absolute error when using the new solution – however, this error decreased with continued practice and was retained in the second baseline (Fig. 2, top). There was a significant main effect of block (F(3.15,148.21) = 23.1, p < .001). Comparisons indicated that absolute error in B1 was lower than N1 (p = .013), absolute error in N4 was lower than N1 (p < .001) and there was no difference between N4 and B2 (p = .341). The improvements in task performance using the new solution were also reflected in a decrease in the number of collisions (t (47) = 6.30, p < .001). (Fig. 2, middle)

**Figure 2.**
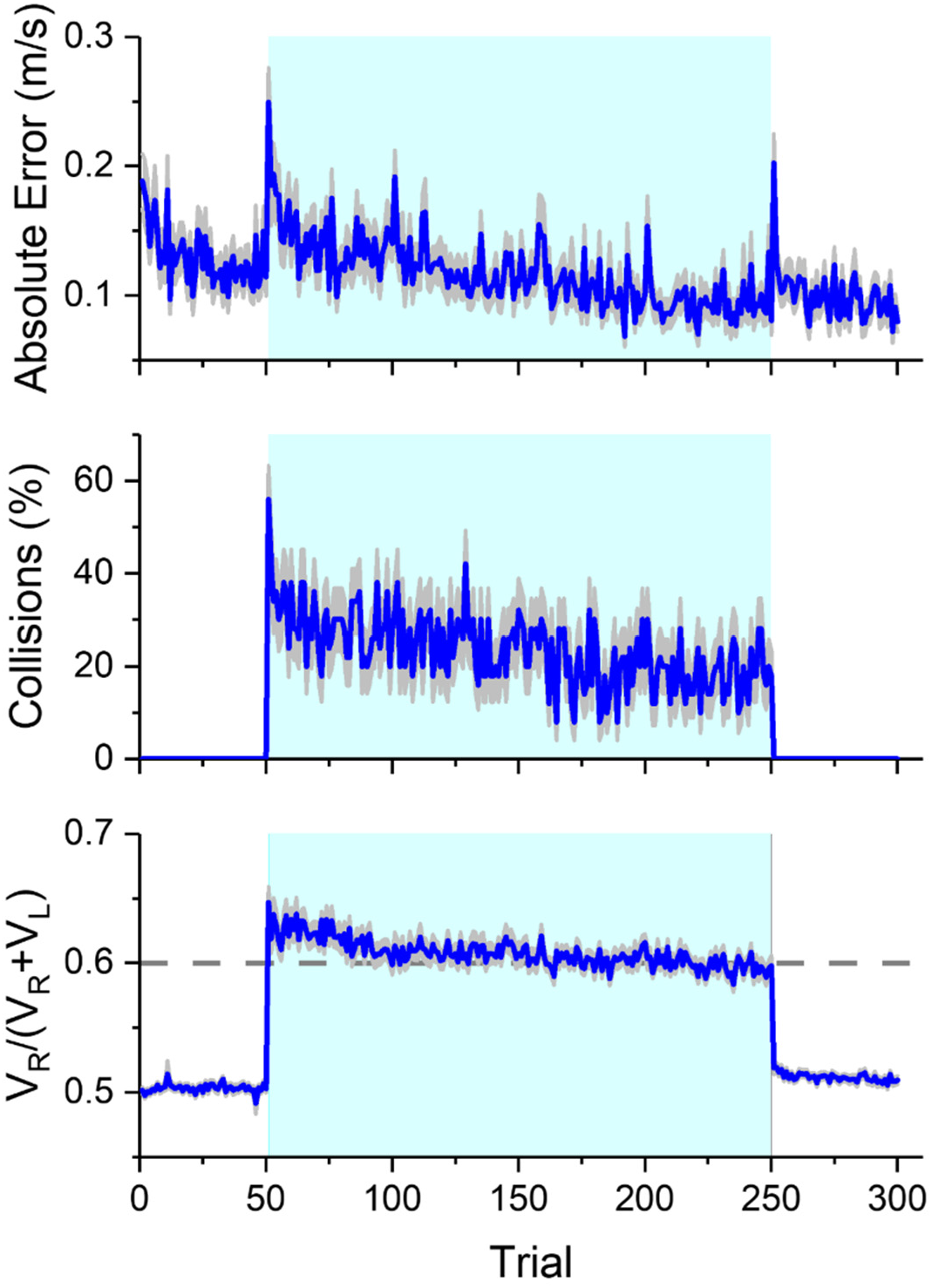
Task performance and coordination during practice. The absolute error, number of collisions, and the contribution of the right hand to the total puck speed are shown as a function of practice. Trials in the novel condition (51-250) are highlighted in blue. Participants were able to adapt to the new solution as indicated by a reduction in absolute error, decrease in the number of collisions, and adjusting the right hand contribution to the ideal level of 0.6. Error bars indicate one standard error (between-participant).

In terms of the coordination, participants showed a change in the speed ratio during the novel blocks but this returned to the original coordination pattern in the second baseline (Fig. 2, bottom). There was a significant main effect of block F(3.04, 143.09) = 334, p < .001). Comparisons indicated that the right hand ratio increased between B1 and N1 (p < .001), decreased from N1 to N4 (p < .001), and then decreased again from N4 to B2 (p < .001).

In terms of task space variability, there was a general decrease in variability across practice (Figs. 3a–3d). There was a significant main effect of block (F(2.83,133.10) = 8.7, p < .001). Comparisons indicated that task space variability (i) was not significantly different between B1 and N1 (p = .790), (ii) decreased from N1 to N4 (p < .001), and (iii) was not significantly different between N4 and B2 (p = .743) (Fig. 3e).

**Figure 3.**
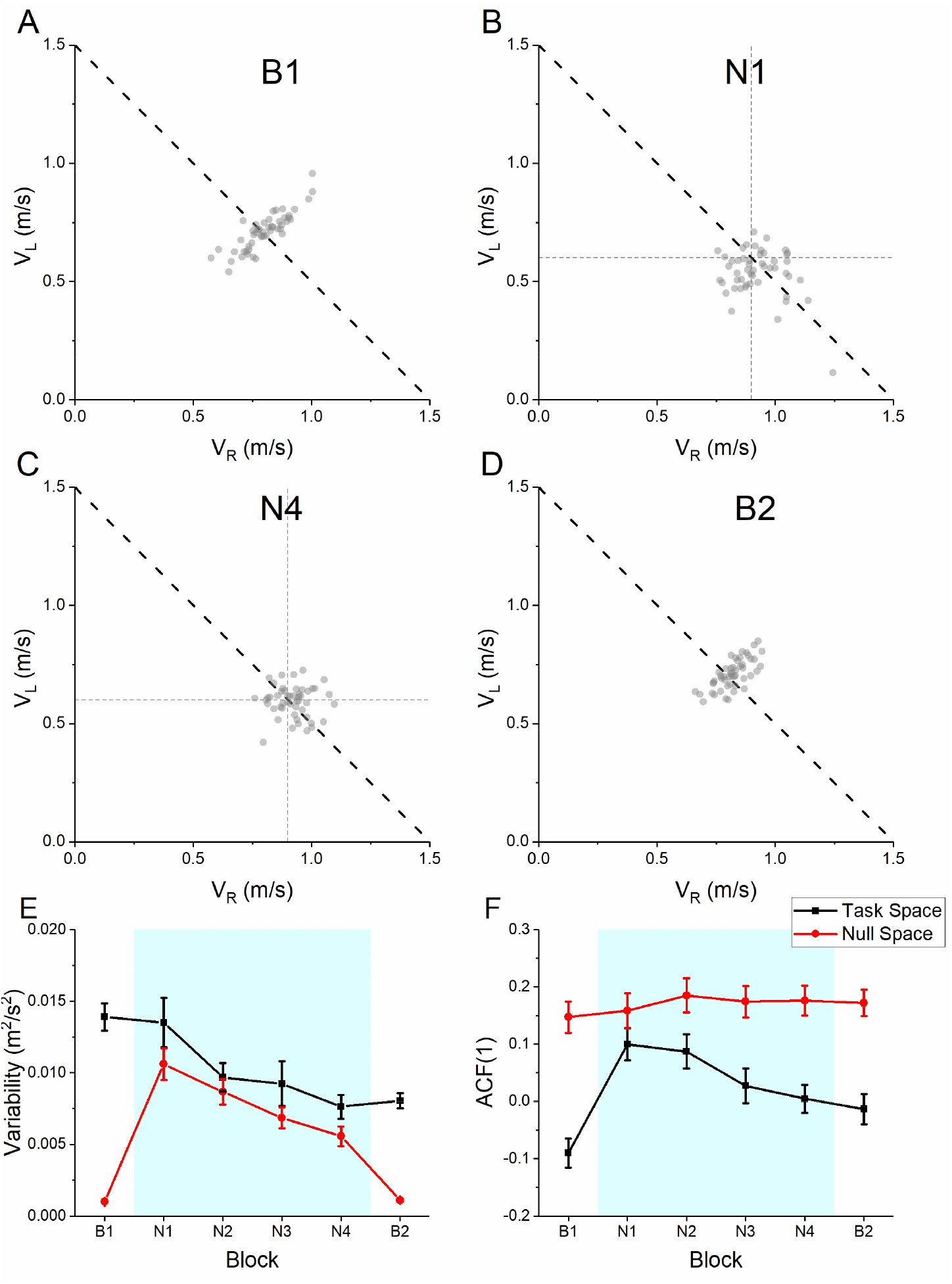
Magnitude and structure of variability with practice. (A)-(D) Sample trials from one participant in block B1, N1, N4 and B2. The solution manifold is represented by the dotted line which indicates the speed combination that lead to a total of 1.5 m/s. The horizontal and vertical reference lines from the axes in (B) and (C) indicate the solution in the novel task. (E) Task and null space variability with practice. Task space variability showed a general reduction with practice. On the other hand, null space variability showed a marked increase in the novel conditions. (F) Autocorrelation. The structure of variability was computed using a lag-1 autocorrelation. In the task space, there were marked increase in the autocorrelation in the novel conditions, but the null space showed no specific trend across practice.

In terms of null space variability, there was an increase in null space variability at the start of novel task, followed by a general decrease until the end of the last novel block, and a sudden decrease when going back to the baseline (Fig. 3e). There was a significant main effect of block (F(2.70,126.88) = 47.3, p <.001). Comparisons indicated that null space variability (i) increased from B1 to N1 (p < .001), (ii) decreased from N1 to N4 (p < .001), and (iii) decreased from N4 to B2 (p < .001).

In terms of the task space autocorrelation, there was an increase in the lag-1 autocorrelation at the start of the novel task, followed by a general decrease until the end of practice (Fig. 3f). There was a significant main effect of block (F (5, 235) = 8.18, p < .001). Comparisons indicated that the autocorrelation (i) increased from B1 to N1 (p < .001), (ii) decreased from N1 to N4 (p = .006), and (iii) was not significantly different between N4 and B2 (p = .604).

In terms of the null space autocorrelation, there was no significant change in the structure during practice (F(5, 235) = .272, p = .928) (Fig. 3f).

### Predicting individual differences

#### Initial learning on the task

Task space variability at baseline was weakly and positively correlated to the initial error at the novel task, indicating that higher variability at baseline resulted in worse initial task performance (Fig. 4a). Initial error at N1 was significantly correlated with task space variability at B1 (r = .398, p = .005) but the correlation with null space variability at B1 (r = .324, p = .025) did not meet the Bonferroni corrected level of significance (Fig. 4b). The difference between these two correlation coefficients was not significant.

**Figure 4.**
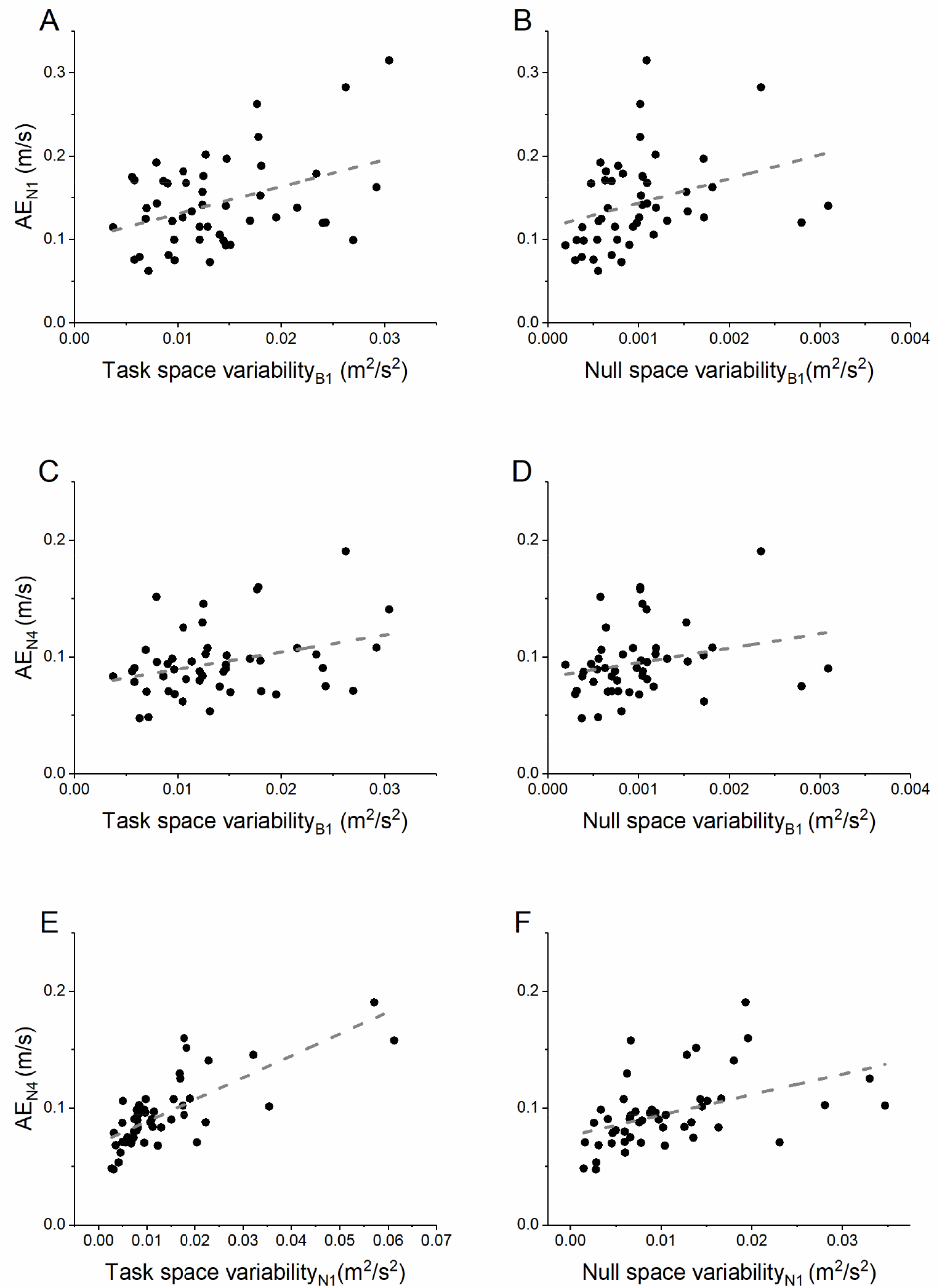
Predicting individual differences in initial and final error from movement variability. (A) Task space variability at baseline was weakly positively correlated with initial error on the novel task (block N1), indicating that individuals showing higher variability had higher initial errors. (B) Null space variability had a nonsignificant correlation with the initial error at N1. When predicting the final error on the novel task (block N4), these correlations were not significant either for movement variability at baseline in either (C) task space or (D) null space. However, a strong positive correlation was observed when correlating the final errors (block N4) to the movement variability in the novel task (block N1) for both the (E) task space and (F) the null space. Once again, though this correlation was positive, indicating that individuals with higher variability at the start tended to have higher errors on the task at the end.

#### Final learning on the task

Task space and null space variabilities at baseline were not correlated with final performance on the novel task (Figs. 4c–4d). The correlation of final errors at N4 with task space variability at B1 (r = .326, p = .024) and with null space variability at B1 (r = .249, p = .088) did not meet the Bonferroni corrected level of significance. Once again, the difference between these two correlation coefficients was not statistically significant.

When we examined if the final performance at the novel task could be predicted by task and null space variabilities during the first block of the novel task (instead of the baseline), we found strong positive correlations, indicating that higher variability in the initial block were associated with higher errors (Figs. 4e–4f). The correlations of final errors at N4 with task space variability at N1 (r = .751, p < .001) and null space variability at N1 (r = .447, p = .001) were significant. The difference between these two correlation coefficients was also significant with a higher correlation for the task space variability (p = .01).

## Discussion

The goal of this study was two-fold – (i) to characterize changes in task performance and movement variability when moving to a new solution to perform the task, and (ii) to examine if movement variability at baseline could predict the ability to perform the task using a new solution. Overall, we found that (i) moving to a new solution resulted in changes in task performance and also in the amount and structure of movement variability, (ii) movement variability at baseline was a poor predictor of the ability to perform the task using a new solution.

### Changes in movement variability when learning a new solution

We found that learning a new solution was associated with several features. From a task performance standpoint, there was a sudden increase in absolute error which was then gradually reduced with practice. This was also associated with an increase in the null space variability (not only in absolute terms but also in relative terms to the task space variability) indicating that participants were exploring the null space to find the new solution. This null space variability decreased with additional practice but was still much higher than what was observed in both the first and second baseline blocks. This effect is particularly surprising because in the novel blocks, the solution space was restricted and therefore the null space variability was not irrelevant to task performance. These suggest that the presence of higher null space variability, by itself, is not always “good” and may potentially reflect the task difficulty (Latash 2010; Scholz et al. 2001) or the use of a solution that is not particularly stable (Ranganathan and Newell 2013).

In addition to the amount of variability, there were also changes in the structure of the variability as measured by the lag-1 autocorrelation function in the task space. The lag-1 autocorrelation in the task space is an index of how errors on the previous trial are being used to correct the next trial, and this was negative in the baseline condition, which is typical for novice performance. However, when moving to the new solution, this lag-1 autocorrelation became positive, indicating that participants likely placed less emphasis on immediately correcting task errors when they were trying to learn the new solution. However, with practice, the lag-1 autocorrelation once again became closer to zero, indicating that participants might be using a learning rate that minimizes the overall variance (van Beers et al. 2013). Surprisingly, we found no effect of the novel task on the autocorrelation in the null space (even though participants had feedback in the null space during the novel task from the left/right motion of the puck), suggesting that the sudden increase in the task space autocorrelation was not due to changes in how participants corrected deviations in the null space.

### Predicting individual differences

Given that a prior study (Wu et al. 2014) had shown that rates of learning were positively correlated with variability, we had expected that initial errors would be smaller for individuals with higher null space variability at baseline (note that in our study, the null space variability is the ‘task relevant’ dimension in the terminology of Wu et al. because this is the intended direction along which exploration should occur to find the new solution). However, we found that the opposite was true – higher null space variability (as well as task space variability) was associated with higher errors, indicating that individuals with greater variability showed lesser ability to produce a consistent outcome using the new solution. This was also seen in the correlations for final performance using the new solution.

We did find a strong ‘specificity’ effect when predicting the final performance at the novel task – movement variability when initially learning the task was highly predictive of final performance using the new solution. But these correlations were once again positive, indicating a detrimental role for variability in learning a new solution. Overall, these results highlight that in the current context, motor variability (both at baseline and the initial learning of the new solution) was more indicative of ‘noise’ in the nervous system – individuals with higher variability showed slower exploration to the new solution and continued to have higher errors in the task. On the other hand, individuals with lower movement variability at baseline were actually ‘more flexible’ – i.e. not only did they perform the task better at baseline, they could also adapt to the new movement solution more easily.

Two issues are critical when considering these findings in the context of prior work – the role of the task, and the measurement of variability. From a task viewpoint, the design of the task and the knowledge of the task goal is an important context modulating the importance of variability in learning. Prior work examining the role of variability have generally focused on adaptation tasks (He et al. 2016; Singh et al. 2016; Wu et al. 2014) or reinforcement-based paradigms using simple tasks (Wu et al. 2014). Adaptation tasks are characterized by adjustments to systematic errors (i.e. changes in constant error or ‘bias’) and several have argued that adaptations to force fields or visuomotor rotations are distinct from tasks where there is an underlying change in the skill (Krakauer and Mazzoni 2011; Sternad 2018). Similarly, in reinforcement learning paradigms, the role of motor variability can be over-estimated because the learning in these tasks is primarily the learning of the task goal, and does not necessarily involve a change in skill. For example in one study (Wu et al. 2014), participants were shown a curve to trace but the actual learning was evaluated on another shape that they were unaware of. This meant that the only way participants could improve on this task was to discover this task goal through trial and error – i.e. identify what the shape of the curve they were being rewarded on. While these prior results advance our knowledge by showing that humans can use motor variability to explore new solutions, in our view, they are less likely to be relevant for many real-life contexts where the task goal is known to the learner in advance. In contrast to these paradigms, in our study, the task goal was known to the learner in advance and learning primarily involved controlling motor variability over multiple trials – we believe this may more closely reflect real-life contexts in motor learning.

The second issue is related to the measurement of variability. A central problem in understanding the role of motor variability is to distinguish ‘noise’ from ‘exploration’ (Therrien et al. 2015). Dhawale and colleagues (Dhawale et al. 2017) attempted to reconcile the somewhat contradictory findings of a metaanalysis on variability (He et al. 2016) by hypothesizing that one critical difference may be related to measuring variability in the presence or absence of feedback. Specifically, measuring variability with task relevant feedback could reflect the ‘noise’ component whereas variability measured without feedback yielded the true ‘exploratory’ component. Our experiment provided a unique test of this hypothesis because the task space variability had feedback, whereas the null space variability in the baseline conditions did not. However, both components of variability showed positive correlations with errors in the task, suggesting that this difference cannot fully account for such discrepancies in the results.

### Conclusion

Overall, these results caution against the use of ‘observed’ motor variability as a predictor of future learning (Ranganathan et al. 2020; Ranganathan and Newell 2010b). The observed variability in a given context is only a ‘snapshot’ of the system’s behavior and may not fully reflect the full potential of the system, which may explain why predictions using variability outside of the specific task context are likely to be less useful. Our results are consistent with conclusion (He et al. 2016) that there is no single relation between variability and learning that generalizes to all contexts, and highlights the need for further work using tasks representative of real-world learning to fully understand the role of variability in motor learning.

## Acknowledgment

We thank Federica Danese for assistance with programming the robot. This work was supported by NSF grant 1823889 to RR and NSF grant 1804053 to CK.

## Author contributions

RR, MH-L and CK conceived and designed research

ML and SC performed experiments

RR, ML, SC and RL analyzed data

RR, MH-L and CK interpreted results of experiments

RR prepared figures

RR drafted manuscript

RR, ML, SC, RL, MH-L and CK edited and revised manuscript

RR, ML, SC, RL, MH-L and CK approved final version of manuscript

